# Fusion-negative rhabdomyosarcoma orthotopic tongue xenografts for study of invasion, intravasation and metastasis in live animals

**DOI:** 10.1101/2023.09.21.558858

**Authors:** Sarah M. Hammoudeh, Yeap Ng, Bih-Rong Wei, Thomas D. Madsen, R. Mark Simpson, Roberto Weigert, Paul A. Randazzo

## Abstract

PAX3/7 Fusion-negative rhabdomyosarcoma (FN-RMS) is a childhood mesodermal lineage malignancy with a poor prognosis for metastatic or relapsed cases. Towards achieving a more complete understanding of advanced FN-RMS, we developed an orthotopic tongue xenograft model for studies of molecular basis of FN-RMS invasion and metastasis. The behavior of FN-RMS cells injected into murine tongue was examined using in vivo bioluminescence imaging, non-invasive intravital microscopy (IVM), and histopathology and compared to the prevailing hindlimb intramuscular and subcutaneous xenografts. FN-RMS cells were retained in the tongue and invaded locally into muscle mysial spaces and vascular lumen. While evidence of hematogenous dissemination to the lungs occurred in tongue and intramuscular xenografts, evidence of local invasion and lymphatic dissemination to lymph nodes only occurred in tongue xenografts. IVM and RNA-seq of tongue xenografts reveal shifts in cellular phenotype and differentiation state in tongue xenografts. IVM also shows homing to blood and lymphatic vessels, lymphatic intravasation, and dynamic membrane protrusions. Based on these findings, the tongue orthotopic xenograft of FN-RMS is a valuable model for tumor progression studies at the tissue, cellular and subcellular levels providing insight into kinetics and molecular bases of tumor invasion and metastasis and, hence, new therapeutic avenues for advanced FN-RMS.

## Introduction

Rhabdomyosarcoma, which accounts for 50% of pediatric soft tissue sarcomas, is a malignancy of the mesodermal lineage most commonly occurring in children and adolescents. This type of sarcoma is evenly stratified into 2 major molecular subtypes: PAX3/7 fusion-positive rhabdomyosarcoma (FP-RMS) and PAX3/7 fusion-negative rhabdomyosarcoma (FN-RMS) (1). Rhabdomyosarcoma cases can also be stratified into 4 major histological subtypes including embryonal (∼60%), alveolar (∼20%), pleomorphic (∼10%), and spindle/sclerosing (∼10%) (2). While FP-RMS is frequently associated with the alveolar subtype (∼80%), embryonal RMS cases are predominantly FN-RMS (>95%) and mainly arise in the head, neck, and genitourinary system (3, 4). Therapeutic advances substantially improved the prognosis of patients with non-metastatic or advanced FN-RMS resulting in 5-year survival rates of 70-80%; however, the prognosis for metastatic or recurrent cases has not changed in the last two decades, with an overall survival rate of ∼30% (5).

Incomplete understanding of the molecular basis for invasion and metastasis of FN-RMS, has hindered rational approaches for the development of therapeutics. Advances are, in part, dependent on experimental models of FN-RMS. Currently available experimental models of FN-RMS, though valuable, by themselves have provided only limited insights into invasion and metastasis. Several approaches have been used to model FN-RMS progression. Genetically engineered animals, including mice (GEMMs) or zebrafish have been the source of advances in understanding tumor behavior, but are limited in that the cell(s) of origin of FN-RMS is yet to be identified (6). Xenograft models based on either patient-derived cell lines or patient-derived tumor xenografts (PDX), most commonly with subcutaneous (7–15) and intramuscular (16–22) injections; however, FN-RMS does not occur in either site in patients, and neither local invasion nor metastasis have been reported in these xenograft model systems. To overcome this limitation, other approaches were explored including tail vein (23–25), intraperitoneal (26–28), and supraorbital injections (29, 30). Although intravenous and intraperitoneal models present with metastases to the lungs, lymph nodes, and organs in the intraperitoneal cavity, these sites are neither physiologically relevant nor do they inform on invasion or how it is linked to metastasis.

Here, we established an orthotopic model based on the injection of patient-derived cell lines into the mouse tongue. The tongue has two advantages over other injection sites. First, primary lesions in FN-RMS have been reported to occur in the tongue (31, 32). Second, the site is ideal for non-invasive intravital microscopy (IVM), a powerful tool to follow the dynamics of the various stages of tumor progression, invasion and metastasis within the same animal for extended period of times (33). We combined live animal imaging techniques and necropsy/histopathology to compare the tongue orthotopic injections to the hindlimb intramuscular and subcutaneous injections. We found that the rate of tumor engraftment was similar among the three sites, but tumor growth was greater in the IM and SC sites. Hematogenous mediated metastasis occurred for both IM and tongue injections, while lymphatic mediated metastases and local invasion were only observed with tongue injections. Using IVM, vascular homing and intravasation into lymphatics in real time was detected supporting hematogenous and lymphatic tumor dissemination. Morphological changes consistent with partial myogenic differentiation was observed in both IVM and in H&E-stained tissue sections and was supported by transcriptomics analysis.

We conclude that engrafting FN-RMS cells into the tongue enables the examination of the invasion and metastasis cascade including (1) cell migration and local invasion, (2) vascular/lymphatic intravasation, and (3) interactions with the tumor microenvironment at the cellular and subcellular levels, which can provide novel insight into cellular and molecular bases of FN-RMS progression in vivo.

## Results

### Characterization of tumor growth in three distinct xenograft sites for FN-RMS cell lines

To set up a suitable orthotopic model system to investigate local invasion and distant metastasis we compared three sites for the engraftment of FN-RMS cells, namely, the tongue, the gastrocnemius muscle, and the flank of immunocompromised mice. To this end, in each site, we injected 2 million FN-RMS cell suspended in a Matrigel and media mixture to promote tumor engraftment, as previously reported (16). We used bioluminescence imaging to characterize the progression and dynamics of the injected cells expressing luc2. For the tongue model we used three cell lines: RD, JR-1, and SMS-CTR; whereas for the other two sites only the RD cells. We monitored tumor progression over a period of 8 weeks followed by euthanasia and necropsy for histopathology.

In the tongue model, cells were retained within the initial 2 weeks (100% of RD and JR-1 xenografts and 80% of CTR xenografts) (Fig. 1A and Supplementary Fig. 1 A,B). Regardless the cell line, tumors progressed beyond 2 weeks with three predominant patterns: (1) semi-linear growth in tumor size, (2) reduction in tumor size 3-4 weeks after injection followed by a growth phase, and (3) continuous reduction in tumor (Fig. 1A and Supplementary Fig. 1 A,B). In 30% of the mice with RD xenografts, we observed early signs of invasion and metastasis to the mandible and locoregional lymph nodes within the initial three weeks after injection (Fig. 1A; red arrows). These invasive/metastatic tumors either grew semi-linearly (2 out of 3 of the mice: m4 and m10) or disappeared (1 out of 3 of the mice: m3) (Supplementary Fig. 1C) in the following weeks potentially due to immune surveillance and clearance. In contrast, tumors in the IM and SC sites grew exponentially over 8 weeks with no detectable tumor outside of the injection sites (Fig. 1B,C). The results of the bioluminescence imaging analysis, however, could not exclude tumors at secondary sites as the signal could be masked by overlying tissues and/or the strength of signal at the primary site.

**Figure 1.**
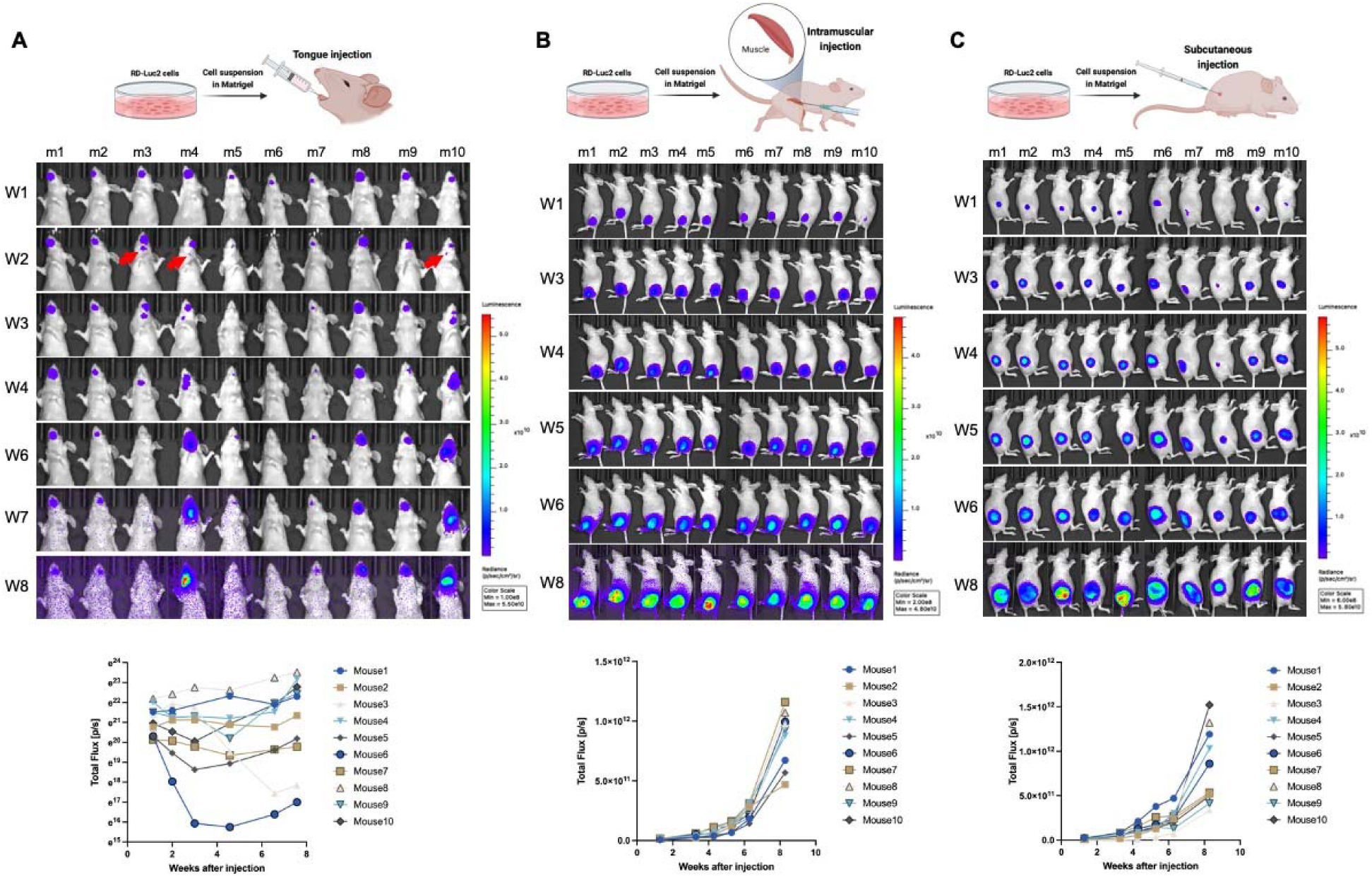
FN-RMS RD cells engraft successfully in the tongue exhibiting different growth dynamics from ectopic xenograft sites such as intramuscular and subcutaneous grafts. Longitudinal bioluminescence *in vivo* imaging for 8-weeks shows tumor progression patterns of RD-Luc2 xenografts injected in (**A**) the tongue, (**B**) hindlimb, and (**C**) subcutaneously. Bioluminescence kinetics presented in the bottom panels of **A** (**, B,** and **C**) show that while hindlimb and subcutaneous xenografts grow uniformly following an exponential growth curve, variations in growth patterns are observed in tongue xenografts. Red arrows in (**A**) point to initial observation of potential invasion/metastasis to the submandibular region (mandible or cervical lymph nodes).

### Histopathology of tumors at the three sites of injection

Hematoxylin and eosin (H&E) stained sections of tongue tissues with injected RD cells revealed poorly circumscribed (Fig. 2A; white dotted box) neoplastic foci of poorly differentiated pleomorphic myoblastic cells (Fig. 2A-D) with heterogenous morphologies including populations of (1) large, atypical cells with abundant eosinophilic cytoplasm and karyomegaly (Fig. 2B; example indicated by white arrows), (2) smaller hyperchromatic round to polygonal neoplastic cells (Fig. 2B; example indicated by yellow arrows), and (3) occasional enlarged cells with strap myocyte morphology and multinucleation in some instances (Fig. 2C; example indicated by white arrow), consistent with limited differentiation towards myocytes. The neoplastic foci in the tongue replaced myocyte bundles and remnant myocytes with hyper-eosinophilic shrunken cytoplasm undergoing degeneration and pressure atrophy were observed in the regions surrounding the tumors. The tongue tumors were interlaced with connective tissue bands (Fig. 2D; yellow arrows) and foci of hyalinized and myxomatous-like matrix material occurs in some masses particularly in packeted tumor foci consistent with extracellular matrix and tumor microenvironment remodeling. The connective tissue bands subdivide the tumor masses with varying degrees of connective tissue condensation to a degree correlating with time-post injection (Supplementary Fig. 2A).

**Figure 2.**
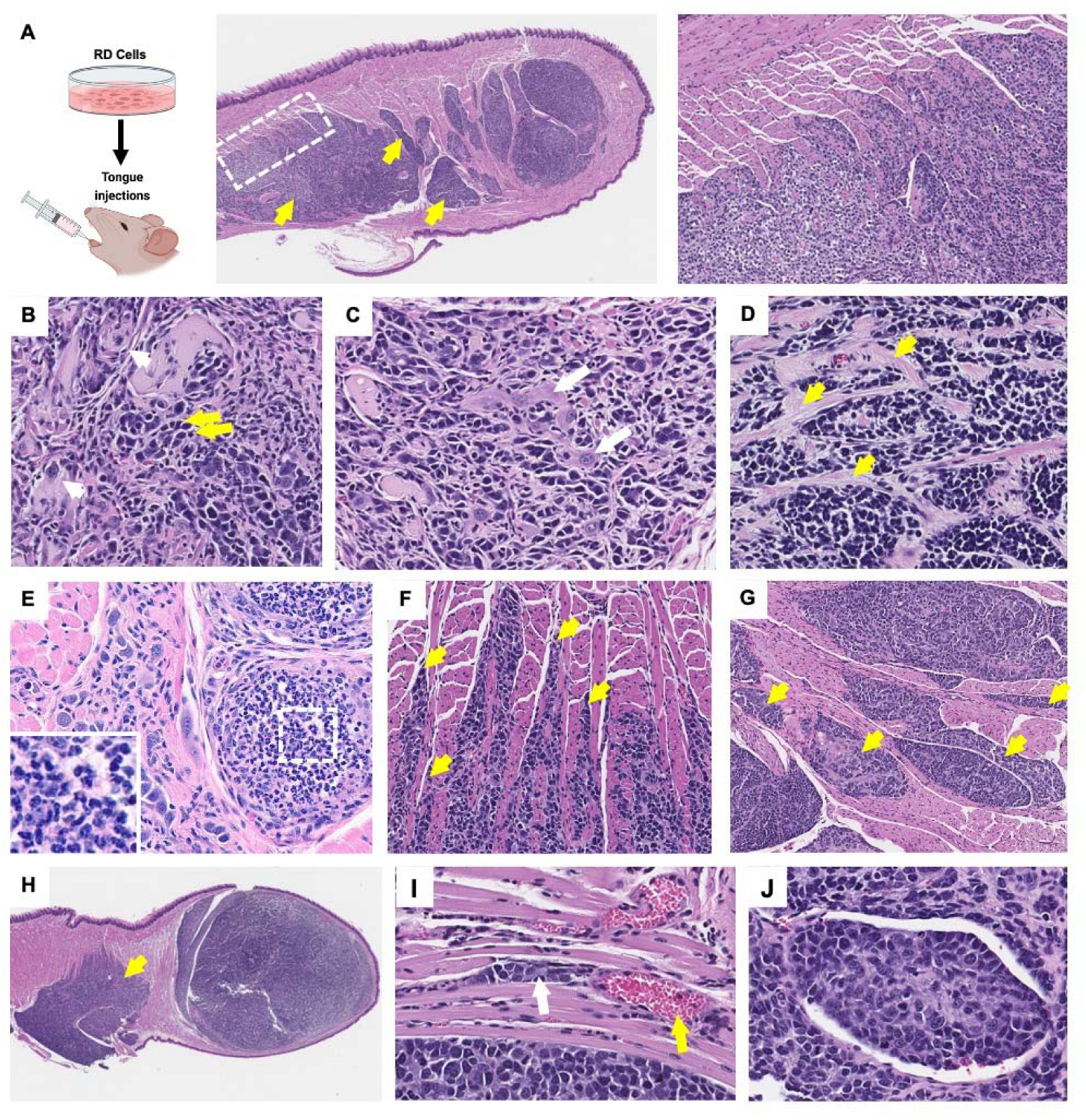
Histopathological analysis shows heterogeneous differentiation states and local invasion in FN-RMS tongue xenografts. (**A**) Histopathology of RD tongue orthotopic xenografts reveals multi-focal poorly circumscribed tumors (dotted box and corresponding zoom-in in the right most panel) invading and forming secondary tumors towards the base of the tongue (yellow arrows). (**B**, **C**) FN-RMS tongue tumors constitute of heterogeneous FN-RMS cell morphologies with different degrees of myogenic differentiation states including large, atypical cells (example in **B**; white arrows), small poorly differentiated round or polygonal cells (example in **B**, yellow arrows), and enlarged multinucleated strap cells (example in **c**, white arrows). (**D**) The tongue tumors are interlaced with connective tissue bands and foci hyalinized and myxomatous-like matrix material (examples in**D**, yellow arrows). **E** (and insets) lymphocytic and monocytic infiltrates are observed in the tongue xenografts. (**F**) FN-RMS cells invade mysial spaces between neighboring muscle cells and fascicles (yellow arrows) (**G**) forming pocketed tumor foci (yellow arrows) and (**H**) secondary tumors deeper in the tongue (yellow arrow). (**I**) Vascular invasion and formation of vascular emboli is also observed in tongue xenografts of FN-RMS cells (white arrow indicates tumor vascular emboli; black * indicates vascular lumen; and red arrow indicates red blood cells).

Lymphocytic and monocytic infiltration was frequently observed in the primary tongue tumors (Fig. 2E; white dotted boxes and corresponding insets at higher magnification), which supports the observed reduction in tumor size at different stages in tumor progression as a potential consequence of immune surveillance.

We also observed a pattern of tumor invasion and growth within the tongue into mysial interphases between myofibers and subdivisions of fascicles (Fig. 2F; yellow arrows). These invasive features resulted in the formation of neoplastic foci in the form of packeted collections in areas replacing myocyte bundles with remnant myocytes (Fig. 2G; yellow arrows). Some of these tumors invaded towards the base of the tongue and grew secondary pocketed foci in these regions (Fig. 2A and H; yellow arrows). Moreover, vascular/lymphatic invasion and intravascular emboli were identified (Fig. 2I).

It was noteworthy that tumor progression differed for the different FN-RMS cell lines in the tongue, as observed by histology (Supplementary Fig. 2B,C). While RD and JR1 xenografts established distinguishable primary tumor foci (Supplementary Fig. 2B), CTR tongue tumors are mostly diminished after 8 weeks post injections with occasional prominent secondary tumors towards the base on the tongue (Supplementary Fig. 2C). These observations indicate that the model can be used to study tumor progression of different FN-RMS cell lines; however, cell-line specific optimization of different parameters (e.g. initial cell concentration, observation window, mouse strain, matrigel content) would be required due to differences in engraftment capacities.

Although the tumors at the IM and SC sites (Fig. 3A,F) were similar to those in the tongue (i.e., both are composed of heterogenous FN-RMS populations including primitive poorly differentiated cells (Fig. 3B,G; white arrows) interlaced with connective tissue bands (Fig. 3B,G; red arrows), there were a number of differences from the tongue. First, the tumors were well-circumscribed clearly separated from surrounding tissues (Fig. 3C,D,H). The exponential growth of the ectopic tumors compressed surrounding tissues and some tumors were engulfed in fibrous or connective tissue fatty layers. Local invasion into skeletal muscle or other neighboring tissue structures was not seen. Second, necrotic cores were observed more frequently in the larger IM and SC tumors compared to tongue tumors (Fig. 3A,F; yellow *). Third, lymphatic intravasation was not detected; however, invasion through vascular walls could be detected in the IM tumors (Fig. 3E). Fourth, there were few or no cytomegalic cells with karyomegaly and strap myocyte morphology compared to the tumors engrafted into the tongue.

**Figure 3.**
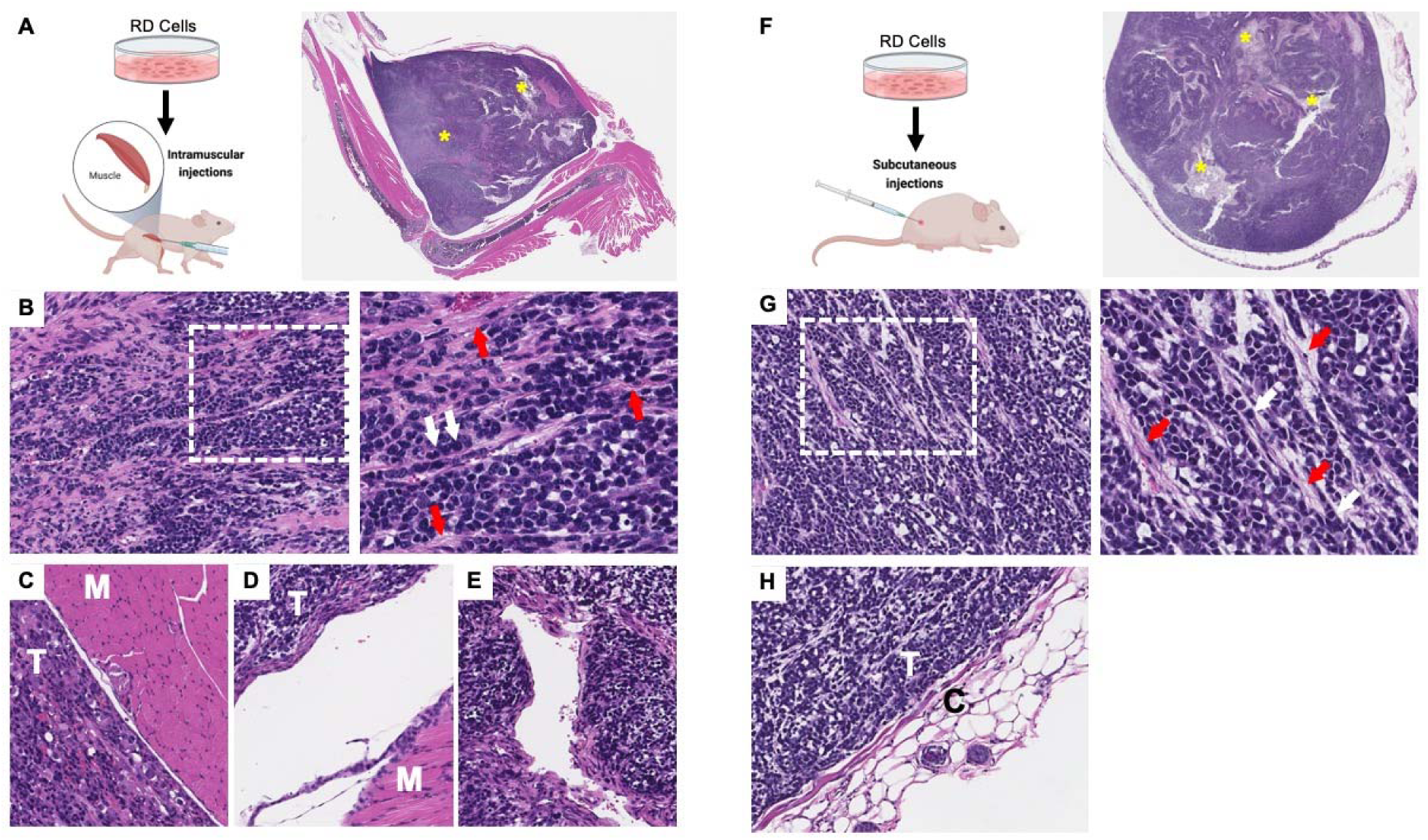
Histopathology of FN-RMS intramuscular and subcutaneous xenograft tumors. (**A**, **F**) Histology of intramuscular injections in the hindlimb and subcutaneous tumors shows well-circumscribed rapidly growing tumors with frequent necrotic cores (yellow *) and consisting of (**B**, **G**; zoom-in panels to the right) predominantly poorly differentiated FN-RMS cells (white arrows) interlaced with connective tissue bands (red arrows). Both (**C**, **D**) intramuscular and (**H**) subcutaneous FN-RMS xenografts were encapsulated and well-separated from surrounding tissues by layers of compressed tissues or connective tissue capsules; T indicates tumor, M indicates skeletal muscle tissue, C indicates connective tissue capsule. While none was observed in subcutaneous xenografts, (**E**) intra-vascular invasion was observed in intramuscular FN-RMS xenografts.

### The metastatic potential of the tumor cells depends on the injection site

To evaluate differences in the rate of metastases among the 3 injection sites at 8-weeks post-injection, we performed a comprehensive necropsy and histopathology of lymph nodes, mandible, salivary glands, lungs, liver, spleen, forelimbs, and hindlimbs.

In the tongue model, tumor cells were detected in cervical lymph nodes, mandibular bone and musculature, salivary glands, and lungs (Fig. 4A-D), which are common metastatic sites in patients (34). In the mandible, poorly differentiated darkly basophilic small mesenchymal pleomorphic rhabdomyosarcoma (RMS) cells with osteoclastic activity and bone lysis were observed (Fig. 4B; white arrow is tumor, yellow arrow is bone matrix). The tumors were densely cellular, forming streams of mesenchymal sarcoma tissue intermingled with bands of connective tissues and remnant myocytes (Fig. 4C). Moreover, intravascular emboli were observed in the mandible suggesting potential intra-/extravasation events through the hematogenous route (Fig. 4D). Tumors were also observed in salivary glands (Fig 4E,F; white arrow is tumor, yellow arrow is salivary gland), although less frequently. Tumors invading the lungs manifested as multifocal lobular pulmonary metastatic RMS (Fig. 4G) with invasion of conducting airways (Fig. 4H) and vascular lumen (Fig. 4I) accompanied by hemorrhage (Fig. 4H, yellow arrows) and lymphocytic infiltration (Fig. 4J, white box and inset). Lung metastases grew in close proximity to vascular structures and, in some cases, tumor cells were found within blood vessels (Fig. 4I). In the lymph nodes, substantial subcapsular accumulation of neoplastic FN-RMS cells, potentially as a consequence of lymphatic dissemination, resulted in replacement of marginal, follicular, and medullary structures of the lymph nodes (Fig. 4K,L; white arrow is tumor, yellow arrow is LN structure).

**Figure 4.**
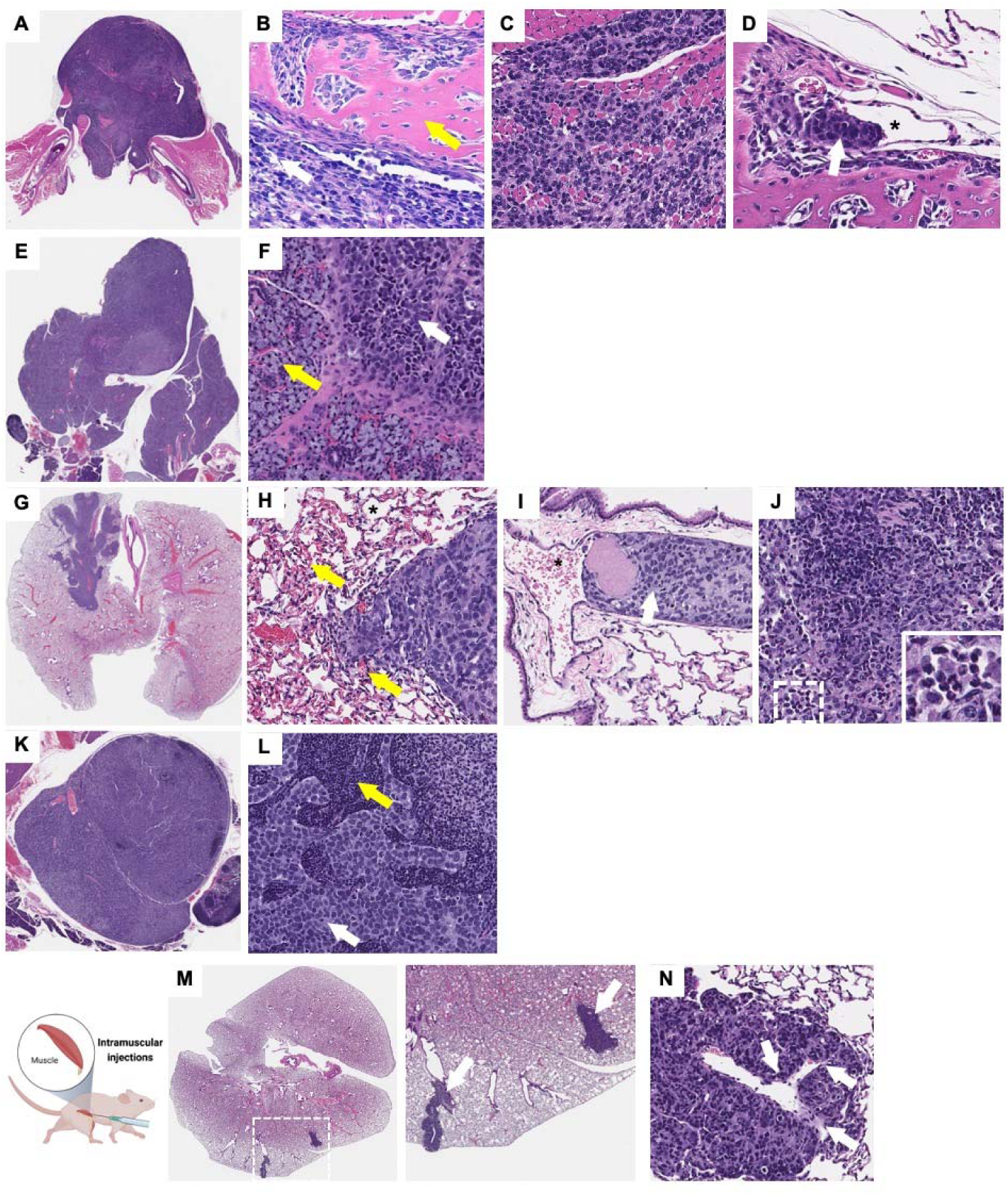
Tongue orthotopic xenografts invade and metastasize to cervical lymph nodes, lungs, lower mandible, and salivary glands while IM xenografts metastasize to lungs. Histopathological analysis shows that RD cells injected into the tongue as an orthotopic site metastasize to (**A-D**) the lower mandible (yellow arrow indicate bone tissue, white arrow indicates tumor), (**E**, **F**) salivary glands (yellow arrow indicate salivary gland tissue, white arrow indicates tumor), (**G**-**J**) lungs, and (**K**, **L**) lymph nodes (yellow arrow indicate LN structure, white arrow indicates tumor). Intravascular invasion can be observed in different sites of metastasis including the (**D**) mandible and (**I**) lungs (white arrows indicate tumor emboli and * indicates vascular lumen). (**M;** higher magnification panel of the right) histopathological analysis shows distant metastasis to the lungs in mice with intramuscular RD xenografts in the hindlimb potentially through hematogenous dissemination as suggested by the intravascular invasion in the lungs (**N;** white arrows indicate invasive tumor regions).

In animals with RD cells injected into the gastrocnemius pelvic limb muscle, metastases to the lungs were observed in ∼40% of mice (Fig. 4M). This was associated with vascular intravasation in the primary tumor (Fig. 3e) and tumor within vasculature in the lungs (Fig. 4N; white arrows representing vascular invasion). Metastases to the lymph nodes were not detected. No metastasis was observed in the mice with subcutaneous FN-RMS xenografts.

In summary, metastasis to the lungs potentially through hematogenous dissemination occurred with both IM and tongue injections, whereas local invasion and metastasis to lymph nodes were only observed when tumor cells were injected into the tongue, as detected by both bioluminescence imaging and histopathology.

### Shifts in the differentiation states are observed in the tongue FN-RMS model

We next examined the transcriptome of tumor biopsies from the tongue RD-EGFP xenografts or associated metastases, which were compared with that of cultured cells. Engraftment of the FN-RMS cells in the tongue results in a transcriptional shift with 2926 upregulated genes and 153 downregulated genes, in comparison to cells in culture. GSEA, functional clustering, and transcriptional factor analysis revealed a prominent upregulation and enrichment of myogenic differentiation/development signatures (e.g. myofibril assembly, striated muscle cell development, intermediate filament organization, sarcomere organization) (Fig. 5A,B; red arrows in a) associated with significant upregulation of the myogenic transcription factors MEF2A and MEF2C downstream targets (Fig. 5C). This enrichment associated with a significant upregulation of myosin heavy chain genes (e.g. *MYH-3, 6, 7, 9*, and *11*), myogenic and associated transcription and enhancer factors (e.g. *MYF5, MYF6, MEF2C, Klf5, NFATC2*), and skeletal muscle structural and dynamics proteins (e.g. *ACTA2, ACTA1, ACTG2, ACTC1, ACTN2, ACTN3, NRAP, TNNT3, TTN*) (Fig 5D).

**Figure 5.**
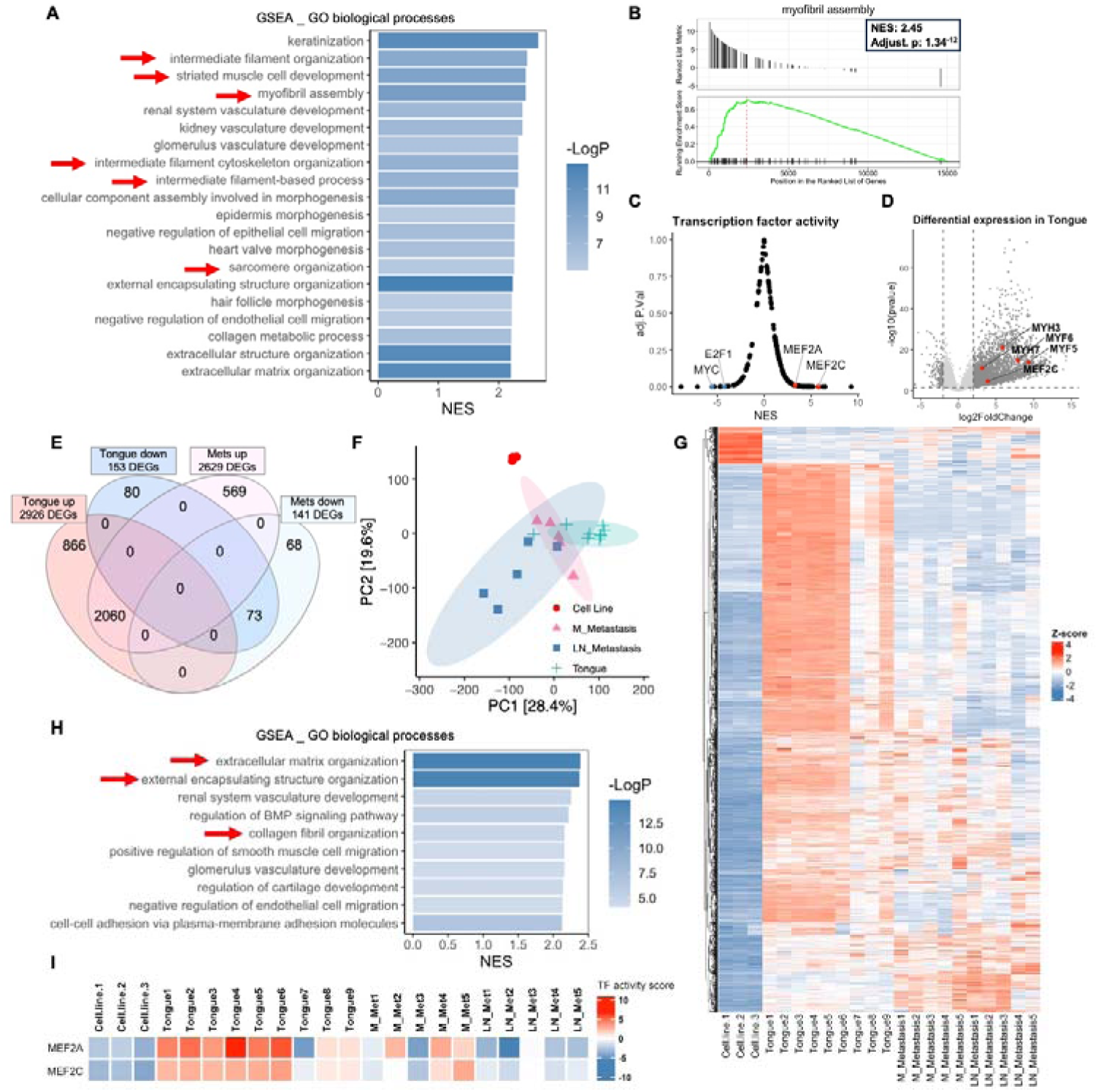
Shifts in differentiation transcriptional signatures between primary and metastatic tumors in the tongue orthotopic xenograft model both compared to cells in culture. (**A**) GSEA analysis of bulk RNA-seq shows that RD tongue orthotopic xenografts are enriched for myogenic differentiation/development and myofibril assembly signatures in comparison to RD-EGFP cells in culture (red arrows). Bar chart representation of the top 20 enriched pathways/biological processes ranked by normalized enrichment score (NES). (**B**) Gene set enrichment (GSEA) plot showing positive, significant enrichment for genes implicated myofibril assembly which are significantly upregulated in RD-EGFP tongue xenograft tumors in comparison to RD-EGFP cells in culture. **C**() Transcription factor (TF) activity analysis performed using Dorothea R package shows a significant positive enrichment for downstream targets of the myogenic differentiation transcription factors MEF2C and MEF2A. (**D**) Volcano plot representation of differentially expression genes (DEGs) in the tongue tumors in comparison to RD-EGFP cells in culture showing examples of DEGs implicated in myogenic differentiation. (**E**) Venn diagram and (**F**) PCA plot and (**G**) heatmap representation of DEGs from tongue tumors show a shift the transcriptional signature between primary tongue tumors and metastatic tumors from cervical lymph nodes and mandible. Venn diagram represents DEGs in the tongue tumors and submandibular metastases (either from the mandible or cervical lymph nodes) using to cells in culture as a reference for both. (**H**) GSEA analysis of the differential transcriptome of submandibular metastases (from the mandible or cervical lymph nodes) shows the loss of the myogenic development/differentiation signatures and the enrichment for extracellular matrix and collagen organization signatures (red arrows) in metastatic tumors in comparison to tongue tumors (**A**); transcriptome of cells in culture was used as a reference. (**I**) TF activity analysis performed using Dorothea R package shows a substantial reduction of MEF2A and MEF2C TF activity in mandibular (M) metastases and to a greater extent in cervical lymph node (LN) metastasis.

While there was substantial overlap in the differential expression signature between the primary and secondary tumors (2060 upregulated genes and 73 downregulated genes) (Fig. 5E,F); discernible transcriptional shifts in mandibular (M) metastases and to a relatively greater extent in cervical lymph node (LN) metastases were detected through principal component analysis (PCA) and hierarchical clustering analysis (fig. 5F,G). Intriguingly, the myogenic differentiation/development signature was lost in the metastatic tumors (Fig. 5H) correlating with the reduction of MEF2C and MEF2A transcription factor activity particularly in LN metastases (Fig. 5I). On the other hand, the differential transcriptome of invasive/metastatic tumors was substantially enriched in genes involved in the organization of the extracellular matrix organization, external encapsulation structures, and collagen fibril signatures genes (e.g. *COL2A1, COL4A5, COL9A3, COL11A1*, and *COL11A2*) (Fig. 5H; red arrows). This finding is consistent with previously published work showing that tumor-draining lymph nodes experience an increase in extracellular matrix remodeling and collagen content (35, 36). These observations support a shift in the myogenic differentiation state towards a less differentiated state in the mandibular and cervical lymph node metastases in comparison to cells in the primary injection site.

### Non-invasive longitudinal imaging of FN-RMS tumor progression in live animals

One of the benefits of using the tongue as an orthotopic site is that it is ideal for non-invasive intravital two photon microscopy (IVM), an approach which enables the visualization of tumor progression within the same animal at cellular and subcellular resolution (33, 37). To perform IVM, we engineered RD cells to stably express cytosolic EGFP. The cells were indistinguishable from the parental cells in terms of morphology and proliferative rate (data not shown). We injected two million cells into the tongue of nude mice and imaged by IVM over a 9-week period. Primary tumors and associated collagen I were easily detected by two-photon microscopy and second harmonic generation (SHG) respectively (38) (Fig. 6A, Movie 1).

**Figure 6.**
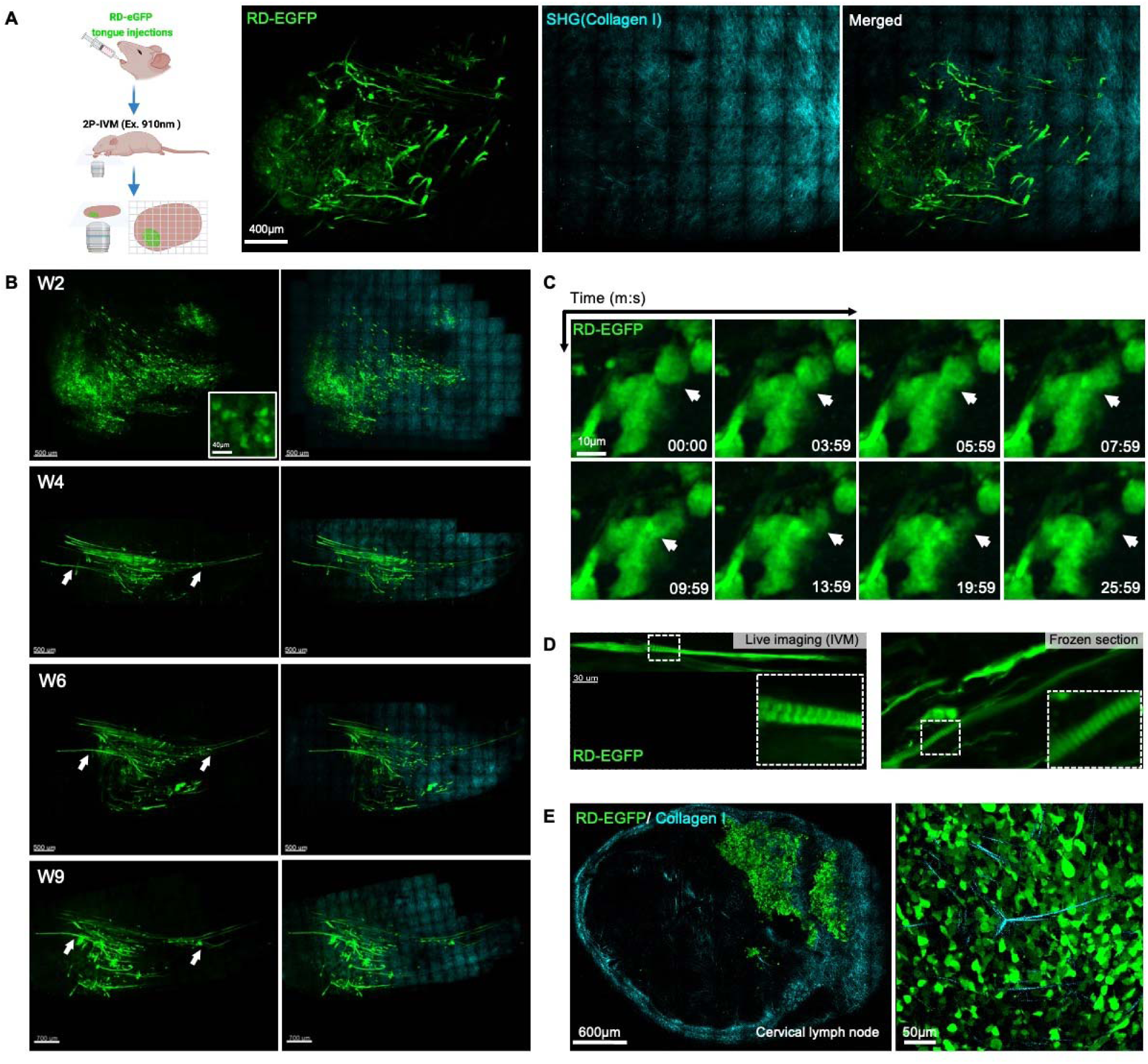
Non-invasive IVM reveals cell fusion and formation of myofibril-like structures in FN-RMS tongue xenografts. (**A**) Two photon intravital imaging of RD cells with constitutive expression of cytosolic EGFP (RD-EGFP; in green) and submucosal layers of collagen I (cyan; detected by second harmonics) after 8-weeks engraftment in the tongue. (**B**) Longitudinal imaging of RD-EGFP tongue xenografts 2, 4, 6, and 9 weeks after cell injection showing the replacement of round individual cell populations observed 2 weeks after injection (W2 and inset) with myofibril-like elongated cell structures which can be observed 4 weeks after injections (W4). These structures persist and grow in length in the following weeks (W4-9). (**C**) Short-term time-lapse two-photon imaging reveals dynamic cell-cell fusion in the RD-EGFP tongue orthotopic xenografts. (**D**) The elongated myofibril-like structures forming in the tongue exhibit signs of myogenic differentiation such as striations extending along their length revealed by two-photon imaging (top panel) and epi-fluorescence imaging of frozen sections (lower panel). (**E**) Two-photon imaging of RD-EGFP cells metastasized to the cervical lymph nodes from the primary tongue xenografts 8 weeks after injection shows the loss of the myofibril-like structures in the metastases; higher magnification image in the panel to the right. Intravital two-photon images in (**A**, **B**, **C**, and **D**) are orthogonal maximum projections of Z-stacks.

As we observed by bioluminescence imaging, within the first two weeks post injection, most engrafted tumors were retained (Fig. 6B). IVM revealed predominantly discrete and individual cells or cell clusters at the sight of injection (Fig. 6B and inset; W2). At later time points, within the field of view with 2-photon imaging, we observed a progressive replacement of individual FN-RMS cells with myofibril-like structures invading towards the base of the tongue (Fig. 6B, arrows; Week4-9). Time-lapse imaging showed cell to cell fusion events as early as 10 days post injections (Fig. 6C, arrows; Movie 2). At later stages (∼5 weeks post injections), striations could be observed throughout these elongated myofibril-like structures, which could also be detected in frozen sections (Fig. 6D and insets). The myofibril-like cell structures grew longer and persisted throughout the period of observation (Fig. 6B). However, these structures were absent in metastases to the cervical lymph nodes, where cells were round or, in some cases, had a mesenchymal morphology (Fig. 6E and zoom-in panel). These observations suggest that a population of the injected FN-RMS cells partially differentiate toward myocytes in the tongue to a greater extent than FN-RMS cells in the lymph nodes, consistent with the transcriptomic analysis. However, we could not find a correlate for these myofibril-like structures in H&E analysis which warrants further investigation.

Using IVM, cell behaviors leading to invasion and metastasis were observed. We used high-resolution time lapse imaging to monitor RD-EGFP cellular dynamics in association with blood and lymphatic vessels (labeled with CD31 eFluor 450-conjugated Antibody and LYVE-1 Alexa Fluor® 594-conjugated Antibody, respectively) (Fig. 7A,B).

**Figure 7.**
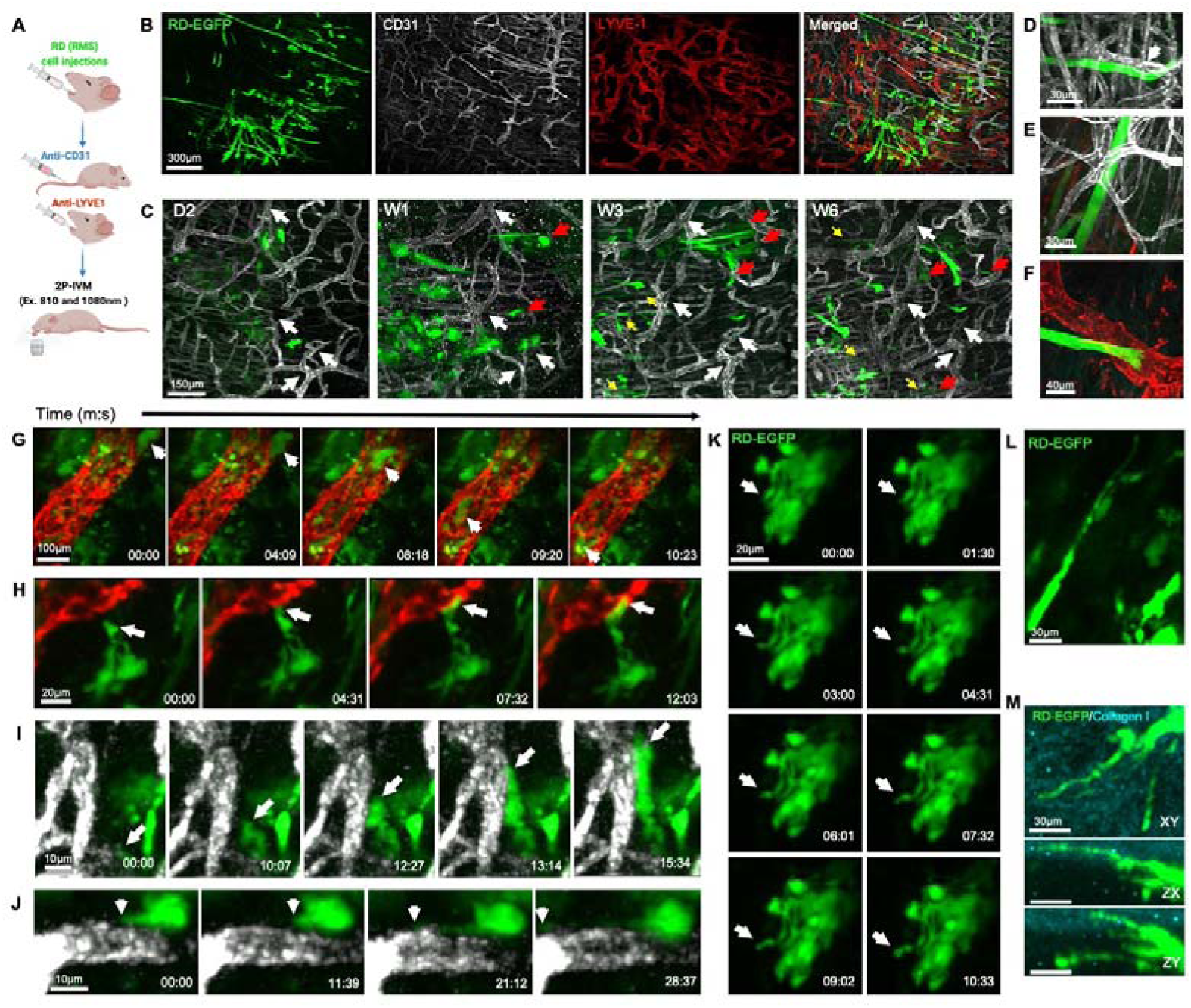
FN-RMS cells home to and intravasate into blood and lymphatic vessels. (**A**) Diagram depicting two-photon intraviral imaging of RD-EGFP cells engrafted in the tongue using CD-31 antibody to visualize blood vessels and LYVE-1 antibody to visualize lymphatic vessels. (**B**) IVM imaging of tongue tumors 4 weeks after injection show the growth of the RD-EGFP cells (green) in close proximity to the abundant blood (white) and lymphatic (red) vasculature in the tongue. (**C**) Longitudinal imaging of peripheral region in the tongue tumor (white arrows indicate vasculature reference points for longitudinal comparison) shows RD-EGFP cells start to seed and migrate into neighboring regions (red arrows) within a week after injections and within 3 weeks elongated myofibril-like structures can be observed arranging partially parallel to blood vessels (yellow arrows in**C** and higher magnification image in **D**). (**E**, **F**) Within 4-9 weeks, the elongated myofibril like structures grow entangled with and impinging on (**E**) blood and (**F**) lymphatic vessels. (**G**) Still images from time-lapse two-photon imaging of the tongue 3 days after injecting RD-EGFP cells depict tumor cells that intravasated into and circulating in lymphatic vessels. The FN-RMS cells migrate and home to (**H**) lymphatic and (**I**) blood vessels where they impinge on the vascular wall and (**J**) form membrane protrusions that trace along the vascular surface. (**K**, **L**) RD-EGFP cells *in vivo* exhibit dynamics formation of long membrane protrusions that extend into surrounding tissues including (**M**) the collagen I rich submucosal layer on the tongue, as shown by two-photon intravital imaging. All intravital two-photon images in this figure are maximum projections of Z-stacks.

Through longitudinal imaging of a region of interest over a period of 6 weeks (Fig. 7C; white arrows indicate reference points), we observed changes in cell organization and localization (Fig. 7C). While initially arranged in clusters of round cells during the first week post injection (Fig. 7C, W1), cells slowly migrated into neighboring regions (Fig. 7C; red arrows) and arranged into chain-like structures partially aligned to blood vessels in the following weeks (Fig. 7C yellow arrows and Fig. 7D). Some FN-RMS myofibril-like structures were entangled with and or in a close proximity to blood vessels (Fig. 7E) and the leading edges of some of these structures projected towards lymphatic vessels (Fig. 7F).

Notably, in the first week after the injection, FN-RMS cells were detected within lymphatic vessels (Fig. 7G, Movie 3) and actively homed in, associated with, or traveled along smaller lymphatic and blood vessels (Fig. 7H,I, Movie 4 and 5), with the formation of cell protrusions that extend along the vascular wall (Fig. 7J, Movie 6). Altogether, these observations further support the histological analysis suggesting invasion and dissemination of FN-RMS cells through the hematogenous and lymphatic routes.

IVM allowed us to observe changes in cellular and membrane dynamics postulated to drive invasion (39, 40). In an *in vivo* environment, tumor cells are surrounded and engulfed by the cellular and acellular elements of the microenvironment resulting in dynamic formation of cell surface protrusions reminiscent of invadopodia. By high resolution IVM time-lapse imaging of tumors within the initial 10 days post injections, we observed dynamic formation of long and thin cell protrusions that invaded collagen I matrix (Fig. 7K,L,M Movie 7). At later stages (e.g. 8 weeks post injections), these events are relatively muted especially for the cells fused into the myofibril like structures.

## Discussion

While prognosis for patients with non-metastatic FN-RMS is relatively favorable with current treatment protocols, the prognosis for patients with metastasis and/or relapsed disease is poor, with little or no advances in treatment in over 2 decades. A rational approach to developing new therapeutics requires an understanding of the molecular and cellular bases of FN-RMS invasion and metastasis, which are still being discovered. Here, we have characterized a xenograft model in which cells derived from human FN-RMS tumors are injected into the mouse tongue as an orthotopic site (31, 32). We found that different than the commonly used hindlimb and subcutaneous xenograft models, xenografts in the tongue orthotopic model invaded adjacent tissues, intravasated into both lymphatics and blood vessels and metastasized to lymph nodes and lung, which are common sites in human disease (34). Furthermore, the tongue model has the advantage of allowing non-invasive intravital imaging, enabling examination of tumor dynamics at the cellular and subcellular levels in live animals in longitudinal studies of tumor-progression. We conclude that the tongue orthotopic xenograft model will be of value in offering insight into the cellular and molecular bases of FN-RMS invasion and metastasis and therefore complement existing models in characterizing the tumor progression cascade*in vivo*.

Different than the widely used intramuscular and subcutaneous xenograft models, the tongue orthotopic xenograft model permits the observation of local invasion at the sight of the primary tumor. Differences in behavior of tumors injected into orthotopic sites from ectopic sites is common. For instance, orthotopic engraftment of colon cancer cells into the cecum of nude mice enhanced invasion into the colonic wall and metastasis to locoregional lymph nodes; which is not observed by the engraftment of other cancer cells (e.g. melanoma cells) in the same location (41). The tongue has been established as an orthotopic xenograft site for head and neck and oral squamous cell carcinomas, with a high primary tumor growth incidence and metastasis to locoregional tumor draining lymph nodes (i.e. cervical lymph nodes) and lungs (42). Similar behaviors are not observed in ectopic xenografts (e.g. subcutaneous xenografts) (43). Here, we found that FN-RMS cells injected into the tongue display similar skeletal muscle invasion dynamics observed in other tumors (e.g. thyroid cancer, Ewing’s bone sarcoma, and oral tongue squamous cell carcinoma), where the tumor cells migrate and align along the mysial interphases between myofibers and fascicles in chain-like multicellular structures, occasionally growing into islands of malignant cells within the skeletal muscle tissues (44–48). In contrast, tumors in the hindlimb or subcutaneous tumors compress surrounding tissue, often with formation of a fibrous capsule circumscribing these tumors in response to the physical compression and extracellular matrix remodeling with no signs of local invasion into neighboring tissues.

Distinct properties of each site we examined might explain the differences in the behavior of the tumors. For instance, the tongue is a unique immune compartment, with specific macrophages resident to the tongue and a high density of blood and lymphatic vessels which enable a rapid response to environmental challenges (49). The vasculature of skeletal muscle, which includes tongue, differs from that in the dermis (50, 51). The extracellular matrix is also distinct in the three injection sites examined in this paper (52). Any or all of these factors could affect the behavior of the injected tumor cells. Plausibly, the differences in rate and routes of metastasis could be explained by differences in density and/or architecture of the blood and lymphatic vessels at the different sites. A high density of lymphatics in the tongue and/or the specific resident macrophages might mediate an enhanced immune response to the developing tumors that could affect tumor behavior. Although athymic nu/nu mice have compromised T-cell mediated humoral immunity, these mice display an elevated NK and macrophages cell-mediated cytotoxicity (53, 54). In the tongue FN-RMS xenograft, unlike the ectopic FN-RMS xenografts, we observe a substantial lymphocytic and monocytic infiltration and inflammation, which can promote invasion through a number of mechanisms (55, 56). The immune response could also explain the difference in growth kinetics from the other sites. While intramuscular and subcutaneous xenografts follow a uniform exponential growth curve, tongue xenografts followed a more nuanced growth pattern with an initial increase in tumor size followed by a diminution and then a third phase which is sometimes an increase in size and, in other cases, resolution of the tumor. It is also possible that differences in the molecular constitution of the microenvironment (e.g. extracellular matrix, growth factors and cytokines) might contribute pro-invasive effects in the tongue, especially considering the proximity to the salivary glands which are reservoirs of growth factors in rodents (57, 58). Regardless of whether there is an effect of rodent specific factors, the model provides a means of studying invasion, determining cell autonomous contributions as well as contributions from the physiologically relevant local environment and inflammatory response.

One advantage of the tongue orthotopic xenograft model is that non-invasive intravital imaging can be used to monitor longitudinal tumor progression and cellular dynamics in a well-defined tumor area. The cell lines used can be genetically manipulated to study different targets of interest. Here, we used cytosolic EGFP labeling of FN-RMS cells to monitor cellular dynamics including general tumor growth, cell arrangement and distribution, morphological changes, formation of membrane protrusions, cell migration, cell-cell fusion, interaction with the microenvironment and vascular intravasation. We observe heterogeneous cellular phenotypes through IVM including primitive rounded cellular morphologies, elongating mesenchymal morphologies and differentiated striated myofibril-like structures in the tongue tumors. This diversity of in cellular phenotypes is reminiscent of the variations in myogenic differentiation states observed in patient FN-RMS tumors (59). However, also revealed by IVM, these heterogenous populations are diminished in cervical lymph node metastases which are dominated by more primitive rounded cells populations. Transcriptomic analysis was consistent with these shifts in differentiation state on engraftment in the tongue and differences between primary and metastatic lesions. Thus, IVM can be used in the tongue orthotopic xenograft model to (1) dissect the contributions of epigenetically distinct population to local invasion, intravasation and metastasis, (2) examine the effect of induced shifts in myogenic differentiation state on response to treatment (60, 61), and (3) examine specific molecular mechanisms driving invasion and intravasation through genetic manipulation.

While the tongue orthotopic xenograft model provides a means of examining molecular mechanisms of local invasion, intravasation and metastasis, limitations of the model include: (1) it cannot provide information about tumor initiation; (2) rodent specific factors might confound interpretation of results, such as the high levels of growth factors secreted from the salivary glands (57, 58); and (3) precise injection technique in the tongue is vital to avoid injuring major blood vessels and injecting the cells into the circulation directly, this latter consideration being applicable to all xenograft models. Nevertheless, the model allows the simultaneous examination of tumor growth, local invasion, cell-cell fusion, migration, and hematogenous and lymphatic dissemination in real time at the cellular and subcellular level and, therefore, an examination of mechanism of invasion, metastasis and the link of the two. Therefore, the model will be a valuable tool for understanding the molecular basis of FN-RMS disease progression and for developing therapeutics to block progression of this disease.

## Materials and Methods

### Stable cell lines generation

The FN-RMS cell lines, RD, JR-1, and CTR, used for the generation of the orthotopic xenograft model in this study were a kind gift from Javed Khan. The RD and CTR cell lines were maintained in High glucose DMEM media and the JR1 cell line with RPMI-1640 media; both containing L-glutamine and supplemented with 10% FBS and 1% penicillin-streptomycin. For *in vivo* live imaging, we generated stable cell lines with constitutive expression of EGFP and fire-fly luciferase 2 (Luc2). EGFP and Luc2 were cloned into a Mammalian Gene Expression Lentiviral Vector backbone under a CMV promoter for stable lentiviral transduction; constructs were generated by VectorBuilder. 1×10^5^ cells were transduced by either vector at Multiplicity Of Infection (MOI) 10 in the presence of 5 μg/ml polybrene for 12 hours. Cells were selected using hygromycin (200μg/ml) 48 hours after transduction for 10 days.

### Animal study

All *in vivo* experiments performed in this study were approved by the National Cancer Institute (NIH) Animal Care and Use Committee (ACUC). Female athymic (nu/nu) nude mice (National Cancer Institute at Frederick), 5-week-old and 17 to 25g, were used in the study. The mice were housed in sterile filter-capped cages, fed and watered ad libitum, in the 12:12 light-dark cycle animal facility.

#### Intramuscular and subcutaneous FN-RMS xenograft models

2 × 10^6^ cells RD cells with stable expression of firefly luciferase 2 (RD-Luc2) were resuspended in Matrigel (Corning; 50% in complete DMEM media) and injected into the caudal thigh muscle of the rear leg or subcutaneously into the flank of 6-7 weeks old female athymic (nu/nu) nude mice. For both models, tumor progression was monitored weekly using Xenogen IVIS Lumina III (PerkinElmer) with Living Image software (PerkinElmer). To measure the luciferase in bioluminescence signal as a correlate of tumor volume, mice were injected intraperitonially with a saturating concentration of D-luciferin (PerkinElmer) based on tumor size (3-6 mg). The bioluminescence signal was captured 20 minutes after injection of luciferin and the tumor burden is represented by the total bioluminescence intensity, photons per second. Mice were imaged weekly for 8 weeks to record tumor growth longitudinally. Necropsy was performed 8 weeks after injections.

#### Orthotopic Tongue FN-RMS xenograft model

The tongue xenograft injections performed as described previously by Amornphimoltham et al. (37). 2 × 10^6^ FN-RMS cells with stable expression of firefly luciferase 2 (RD-Luc2, JR1-Luc2, and CTR-Luc2) were injected into the lateral anterior side of the tongue avoiding main blood vessels. Cells were prepared for a total volume of 20ul per injection in a mixture of 50% Matrigel and 50% complete media to promote tumor engraftment. Animals were fed with soft dough diet on the day of injection. Tumor progression was monitored weekly using Xenogen IVIS Lumina III (PerkinElmer) with Living Image software (PerkinElmer). To measure the luciferase in bioluminescence signal as a correlate of tumor volume, mice were injected intraperitonially with a saturating concentration of D-luciferin (PerkinElmer) based on tumor size 3 mg. The bioluminescence signal was captured 20 minutes after injection of luciferin and the tumor burden is represented by the total bioluminescence intensity, photons per second. Mice were imaged weekly for 8 weeks to record tumor growth longitudinally. Necropsy was performed 2, 4, and 8 weeks after injections.

### Intravital two-photon imaging of RD-EGFP tongue xenografts

For intravital imaging, RD cells with stable expression of EGFP (RD-EGFP) were used for the tongue xenografts. 2 × 10^6^ RD-EGFP cells were injected into the lateral anterior side of the tongue avoiding main blood vessels. Animals were fed with soft dough diet on the day of injection. Tumor growth was monitored using handheld fluorescence flashlight (Nightsea Xite-RB, EMS). Tumor bearing animals were selected within the first week after injection to monitor cell dynamics (5-10 days after injection) and to longitudinally monitor tumor progression every 2 weeks.

For intravital imaging, the mice were initially anesthetized by brief exposure to isoflurane followed by a subcutaneous injection of a mixture of 100 mg/kg and 10 mg/kg xylazine (Anased LA, Vet ONE). The tongue was then gently pulled out using blunt forceps and stabilized with cotton tips on the 37°C heated stage of a two-photon microscope. The tongue hydration was maintained by applying a thin layer of carbomer 940 gel while the eyes were hydrated by a piece of gauze wetted with distilled water. The mice were covered with a gauze throughout the experiment to maintain physiological temperature. Imaging was performed by using an inverted TCS SP8 Dive Spectral Microscope (Leica) equipped with Mai-Tai and Insight X3 tunable lasers (Spectral Physics) and a 37 °C preheated 40X objective (NA 1.10, HC PL IRAPO, Leica). The specimens were excited at 910 nm.

Collagen I (Second Harmonic Generation) was detected by HyD-RLD1 at the emission 447-462nm and EGFP was detected with HyD-RLD2 at the emission 512-549 nm. The tile images of the tumor were collected by bidirectional line scan with 2 line averaging at 600 Hz (320 x 320 pixel; 12 bits per pixel) on XY, and 75 to 100 Z-steps (2 um step size) and stitched using LAS X Navigator. The 3D images with timelapse were acquired by bidirectional line scanning with 2 line averaging at 400 Hz (512 x 512 pixel; 12 bits per pixel) on XY, and 19 to 55 Z-stacks (2 um step size) using Leica LAS X software. All the images were stored as LIF files and processed further using Imaris (Bitplane).

### Intravital two-photon imaging of tumor vascular interactions

To visualize the blood vessels in the tongue, 10ug of eFluor™ 450-conjugated CD31 (PECAM-1) Monoclonal Antibody (Clone 390, 50ul of original concentration 0.2mg/ml, Invitrogen, Cat# 48-0311-82) was injected into the tail vein 20 minutes before imaging. Lymphatics were visualized by injecting 0.4ug of Mouse LYVE-1 Alexa Fluor® 594-conjugated Antibody (20ul of concentration 20ug/ml, diluted 1:10 in normal saline; cat# FAB2125T; R&D systems) into the tongue in a region near the tumor 20 minutes before imaging. The tongue was excited at 810 nm and 1080 nm simultaneously. Collagen I (Second Harmonic Generation) was detected by HyD-RLD1 at the emission 400-410 nm, anti-CD31 was detected with HyD-RLD2 at the emission 443-462 nm, EGFP was detected with HyD-RLD3 at the emission 516-537 nm, and LYVE-1 was detected with HyD-RLD4 at the emission 601-623 nm. The tile images of the tumor were collected by bidirectional line scan with 2 line averaging at 600 Hz (320 x 320 pixel; 12 bits per pixel) on XY, and 75 to 100 Z-steps (2 um step size) using LAS X Navigator and stored as LIF files. 3-D stitched images were constructed using Leica LAS X software. Tumor bearing animals were selected within the first week after injection to monitor tumor-vascular interactions (3-7 days after injection) and to longitudinally monitor tumor cell distribution relative to the vascular plexus (day2, week1, week4, and week6).

### Histological analysis of formalin fixed paraffin embedded (FFPE) and fixed frozen sections

Comprehensive necropsy of primary tumor xenografts (tongue, intramuscular, and subcutaneous tumors) and potential sites of metastasis including lymph nodes (cervical, axillary, brachial, inguinal, and popliteal), mandible, salivary glands, lungs, liver, spleen, forelimbs, and hindlimbs was performed. For formalin-fixed paraffin embedded tissues (FFPEs), The samples were fixed in 4% PFA for 4-5 days, then stored in 70% ethanol at 4 °C and processed Tissue-Tek® VIP5™ automated tissue processor and embedded into paraffin wax. 5 μm tissue sections were mounted on positively charged slides for standard H&E staining using Tissue-Tek®Prisma™ automatic stainer and histopathological analysis.

For fresh frozen, mice were perfused with 10mL 4% PFA in 200mM HEPES pH 7.3 through cardiac circulation and tongues were extracted and immersion fixed for 1h in 4% PFA in 200mM HEPES pH 7.3. Subsequently, the tongues were incubated in a two-step sucrose gradient at 15% followed by 30% sucrose in PBS at 4 °C until tissue sank to the bottom of the container (∼2 hours for 15% sucrose and overnight incubation for 30% sucrose). The sucrose-infiltrated tongues were embedded in O.C.T compound embedding medium (Tissue-Tek) by freezing against a positively charged histology slide on powdered dry ice. 10 μm tissue sections were mounted on positively charged slides for epifluorescence imaging of RD-EGFP tumors using Keyence BZ-X710 Fluorescence Microscope.

### RNA extraction and bulk RNA-seq of primary and secondary tumors

Tumor punch-biopsies were collected from the tongue and metastases in the mandible and lymph nodes in Precellys® CKMix Tissue Homogenizing Kit tubes (Bertin technologies; P000918-LYSK0-A) and snap-frozen in liquid nitrogen and stored at −80 °C until extraction. RNA was extracted from cell pellets using RNeasy Plus Mini Kit (Qiagen; 74134), as per the manufacturer’s instructions. 600μl of the supplied Lysis buffer containing freshly added 2-mercaptoethanol was added per sample. The samples were homogenized using Precellys 24 tissue homogenizer (Bertin technologies) at 6200 RPM for 3 cycles, 30 seconds a cycle. The samples were passed through QIAshredder columns (Qiagen; 79654) to eliminate potential tissue aggregates. Genomic DNA content was eliminated by passing the samples through the gDNA Eliminator spin columns. 600μl of 70% ethanol was added to the tissue lysates, mixed thoroughly and transferred to the RNAeasy spin columns. Following the washing steps with RW1 and RPE buffers recommended by the kit, the samples were eluted in 30ul of RNase-free water. RNA content was quantified using NanoDrop ONE ^c^ (Thermo Scientific) and 100ng from each sample was used for Quality control using Tape Station RNA Regular Sensitivity assay.

We used 800 ng of total RNA samples to generate the library (for both ribodepleted and mRNA). RNA-seq libraries were prepared using the NEBNext rRNA Depletion Kit v2 (Human/ mouse) (NEB #E7405), NEBNext Ultra II Directional RNA Library Prep Kit for Illumina (NEB #E7760) and NEBNext® Multiplex Oligos for Illumina® (96 Unique Dual Index Primer Pairs, NEB, E6440) as per manufacturer’s recommendation. Sequencing was carried out in the Illumina NextSeq 2000 instrument with 101x101 pair end configuration with 8x8 index.

### Bulk RNA-seq bioinformatics analysis

Raw sequencing reads were trimmed for adapters and low-quality bases using Trimmomatic before alignment with the reference genome (Human – hg38). Then the analysis was done in the Partek Flow installed in NIH Helix. The trimmed reads were aligned with hg38 using STAR 2.7.8 and quantified based on hg38_ensembl_release107_v2 annotation model. Differentially expressed genes (DEGs) were identified using the DESeq2 R package (62) and defined as having an adjusted p-value <0.05 and log2 fold change (FC) >2 or <-2. Functional implications of the differentiational transcriptome were explored using Gene Set Enrichment Analysis gseGO() function of the clusterProfiler R package. Differential expression data and functional clustering analysis were visualized using the R packages: ComplexHeatmap() and ggplot2(). Prediction of shifts in transcription factor (TF) activity was analyzed using the R package DoRothEA, which creates a gene regulatory network of TF-target interactions using curated sets of TF and transcriptional targets (63). Overlap between genes differentially expressed in the tongue and metastatic tumors was examined using InteractiVenn (64).

## Supporting information

Move 2

## Acknowledgements

We thank the Frederick National Laboratory for Cancer Research (FNLCR) Molecular Histopathology Laboratory (MHL) for processing tissues for histopathology. We thank the National Cancer Institute Laboratory Animal Sciences Program (LASP) for their technical support and animal care. We thank the CCR genomics core for their support in sequencing the RNA samples and Haiyan Lei and John Shern for alignment of the RNAseq reads. We thank John Shern for insightful discussions and Marielle Yohe for insightful discussions at the initiation of the project. This work was supported by the Intramural Research Program of the National Cancer Institute, National Institutes of Health, Department of Health and Human Services (project number BC007365 to P. A. R., project number ZIA BC 011682 to R. W. and the CCR IVM Core funding number ZIC BC 012030).

## Supplementary Figures

**Supplementary Figure 1.**
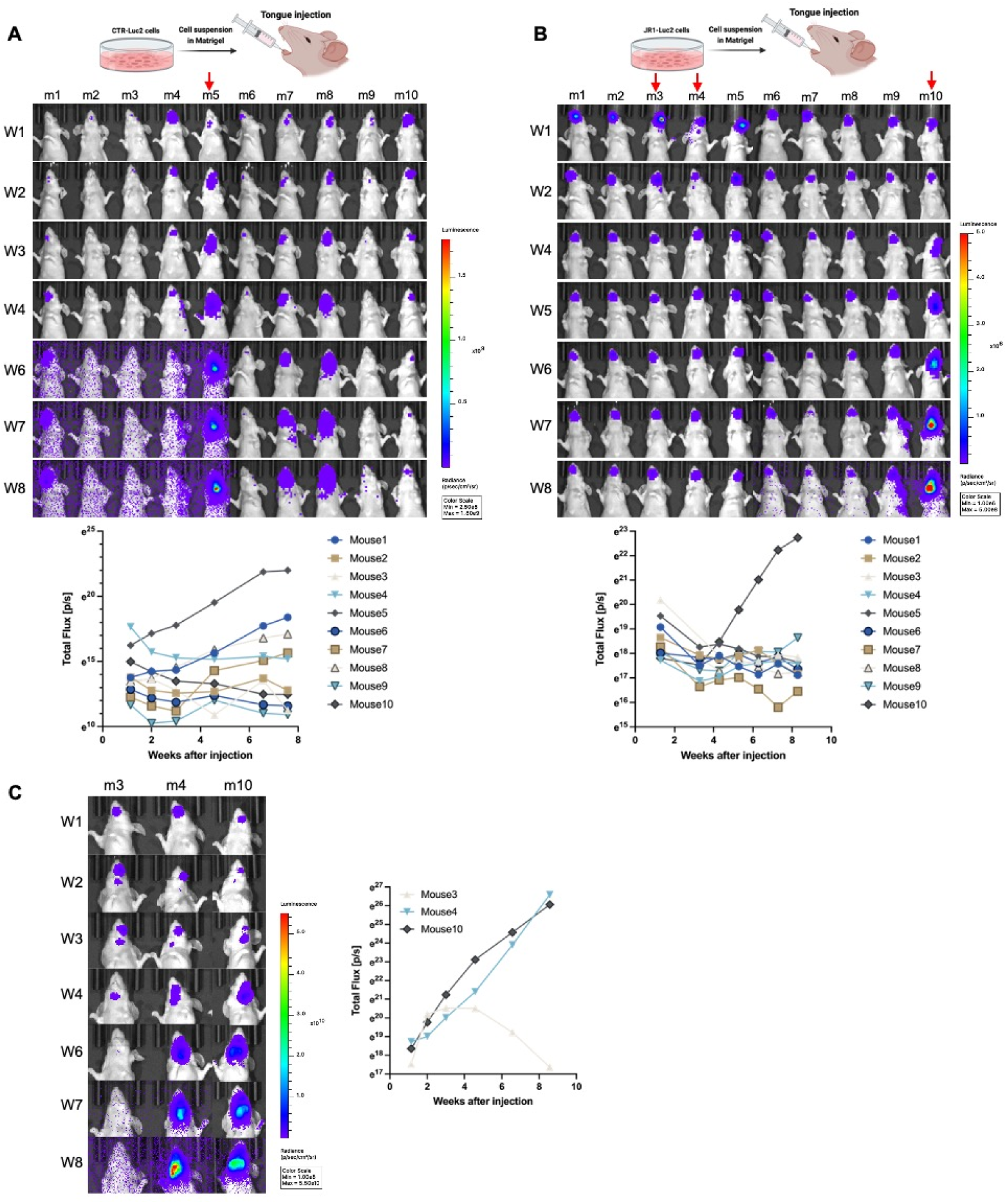
Bioluminescence in vivo imaging to monitor progression of FN-RMS tongue tumors. (**A, B**) Longitudinal bioluminescence imaging to characterize tumor progression of (**A**) CTR-Luc2 and (**B**) JR1-Luc2 xenografts injected in the tongue showing tumor progression. Bioluminescence kinetics are presented in the bottom panels of (**A**) and (**B**). (**C**) Longitudinal bioluminescence imaging shows invasion to the lower mandible or cervical lymph nodes that either progressed semi-linearly (m4 and m10) or resolved during the observation period (m3). Bioluminescence kinetics are presented in the panel of the right of (**C**).

**Supplementary Figure 2.**
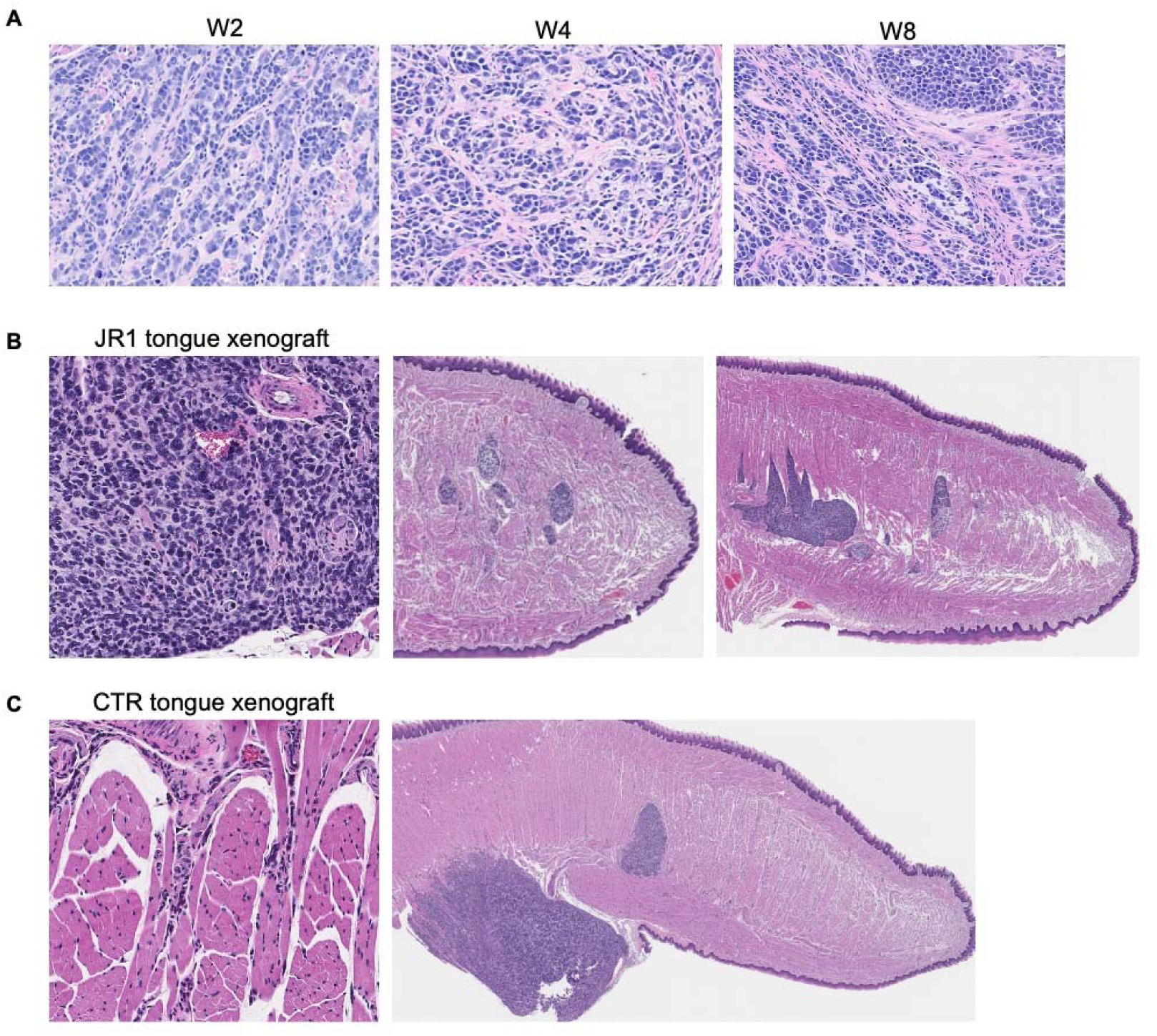
Histology of FN-RMS tumor progression in the tongue. (**A**) Longitudinal histopathological analysis of RD tongue orthotopic xenografts shows variations in connective tissue condensation correlating with time post injection. (**B**) Histology of JR1 tongue tumors showing zoom-in view in the panel to the left, established primary xenograft at the sight of the injection in the middle panel, and an example of occasional secondary tumors in the right most panel. (**C**) Histology of CTR tongue tumors showing zoom-in view in the panel to the left and an example of occasional secondary tumors in the panel to the right.

## Movie Legends

**Movie 1. 2P-IVM of the RD-EGFP tongue tumor (green) 8 weeks after injection.** The video is a 360-rotation of the 3D rendering. Collagen I (cyan) is detected by second harmonics.

**Movie 2. 2P-IVM of cell-to-cell fusion of RD-EGFP (green) in the tongue tumor.** Video is a maximum-intensity projection of an image stack acquired for ∼24 minutes (min:s) with a frame rate of 60 frames/s.

**Movie 3. 2P-IVM of RD-EGFP cells (green) circulating in lymphatic vessels (red) 3 days after injection.** Video is a maximum-intensity projection of an image stack acquired for ∼24 minutes (min:s) with a frame rate of 70 frames/s. Lymphatic vessels were labeled with anti-LYVE-1.

**Movie 4. 2P-IVM of RD-EGFP cells (green) homing to lymphatic vessels (red).** Video is a maximum-intensity projection of an image stack acquired for ∼10 minutes (min:s) with a frame rate of 40 frames/s. Lymphatic vessels were labeled with anti-LYVE-1.

**Movie 5. 2P-IVM of RD-EGFP cells (green) homing to blood vessels (white).** Video is a maximum-intensity projection of an image stack acquired for ∼23 minutes (min:s) with a frame rate of 70 frames/s. blood vessels were labeled with anti-CD31.

**Movie 6. 2P-IVM of RD-EGFP cells (green) extending an invadopodia-like protrusion along the blood vessel (white) wall.** Video is a maximum-intensity projection of an image stack acquired for ∼13 minutes (min:s) with a frame rate of 100 frames/s. blood vessels were labeled with anti-CD31.

**Movie 7. 2P-IVM dynamic membrane protrusion formation in RD-EGFP cells (green).** Video is a maximum-intensity projection of an image stack acquired for ∼10 minutes (min:s) with a frame rate of 35 frames/s.

